# Thalamocortical interactions reflecting the intensity of flicker light-induced visual hallucinatory phenomena

**DOI:** 10.1101/2024.04.30.591812

**Authors:** Ioanna A. Amaya, Till Nierhaus, Timo T. Schmidt

## Abstract

The thalamus has a critical role in the orchestration of cortical activity. Aberrant thalamocortical connectivity occurs together with visual hallucinations in various pathologies and drug-induced states, highlighting the need to better understand how thalamocortical interactions may contribute to hallucinatory phenomena. However, concurring symptoms and physiological changes that occur during psychopathologies and pharmacological interventions make it difficult to distil the specific neural correlates of hallucinatory experiences. Flicker light stimulation (FLS) at 10 Hz reliably and selectively induces transient visual hallucinations in healthy participants. Arrhythmic flicker elicits fewer hallucinatory effects while delivering equal amounts of visual stimulation, together facilitating a well-controlled experimental setup to investigate the neural correlates of visual hallucinations driven by flicker rhythmicity. In this study, we implemented rhythmic and arrhythmic FLS during fMRI scanning to test the elicited changes in cortical activation and thalamocortical functional connectivity. We found that rhythmic FLS elicited stronger activation in higher-order visual cortices compared to arrhythmic control. Consistently, we found that rhythmic flicker selectively increased connectivity between ventroanterior thalamic nuclei and higher-order visual cortices compared to arrhythmic control, which was also found be positively associated with the subjective intensity of visual hallucinatory effects. As these thalamic and cortical areas do not receive primary visual inputs, it suggests that the thalamocortical connectivity changes relate to a higher-order function of the thalamus, such as in the coordination of cortical activity. In sum, we present novel evidence for the role of specific thalamocortical interactions with ventroanterior nuclei within visual hallucinatory experiences. Importantly, this can inform future clinical research into the mechanistic underpinnings of pathologic hallucinations.

## Introduction

The thalamus is a subcortical structure that has diverse functions in filtering sensory information, regulating cortical excitability and integrating information across cortical networks (Sherman and Guillery, 2006; Hwang et al., 2017; Halassa and Sherman, 2019; Shine et al., 2023). The various thalamic functions are served by different thalamic nuclei, whereby first-order nuclei (e.g., lateral geniculate nuclei; LGN) mainly relay and filter sensory information and higher-order nuclei (e.g., ventroanterior, reticular nuclei) are primarily involved in the coordination of activity and information availability across cortices. Thalamocortical hyperconnectivity has been observed in a range of pathologies and altered states of consciousness, such as psychosis (Anticevic et al., 2014a, 2014b), Charles Bonnet Syndrome (Ffytche, 2008), and during psychedelic experiences (Müller et al., 2018; Preller et al., 2019). In some cases, thalamocortical hyperconnectivity has been correlated to the intensity of hallucinatory experiences (Anticevic et al., 2014b; Müller et al., 2018). However, there is not yet a consensus on which thalamic nuclei display changes in connectivity. A better understanding of which thalamic nuclei become hyperconnected can help to unravel the functional relevance of thalamocortical hyperconnectivity during visual hallucinatory experiences.

To investigate which thalamic nuclei display thalamocortical hyperconnectivity, it is crucial to identify an experimental tool that can selectively elicit hallucinatory experiences without additional confounding effects. For example, while thalamocortical connectivity during psychotic states and psychedelic experiences has been extensively investigated (Anticevic et al., 2014a, 2014b; Müller et al., 2017; Ferri et al., 2018; Preller et al., 2019; Avram et al., 2020, 2022; Bedford et al., 2023; Pizzi et al., 2023), hallucinatory experiences within these populations co-occur with other physiological effects, such as long-term compensatory changes for psychotic patients (Lieslehto et al., 2021) and widespread systemic effects arising from pharmacological intervention (Schmid et al., 2015; Luppi et al., 2021; Hirschfeld et al., 2023). This means that the described neural correlates are not necessarily specific to the visual experience. On the contrary, flicker light stimulation (FLS) can reliably and selectively induce visual hallucinatory effects in healthy participants via closed-eye application of stroboscopic light. The flicker-induced effects are mainly constituted of elementary visual hallucinations, referring to the perception of geometric patterns and colours devoid of semantic content (Bartossek et al., 2021; Amaya et al., 2023a). These elementary phenomena are also frequently reported during psychedelic experiences (Schmidt and Berkemeyer, 2018; Prugger et al., 2022), Charles Bonnet Syndrome (Jan and Castillo, 2012) and for some patients with psychosis (Ommen et al., 2019). Therefore, utilising FLS allows the assessment of specific neural effects occurring during visual hallucinations that are also experienced in other pathologies and pharmacologically induced states.

Flicker rhythmicity and frequency interact to produce different intensities of hallucinatory effects (Amaya et al., 2023a) with rhythmic 10 Hz FLS eliciting the strongest visual effects. Recent research identified increased connectivity between anterior, ventral and mediodorsal thalamic nuclei and visual cortices during 10 Hz FLS compared to 3 Hz FLS (Amaya et al., 2023b). However, comparing FLS-induced neural effects across flicker frequencies introduces differences in the amount of visual stimulation. Therefore, we recently developed a frequency-matched arrhythmic flicker that eradicates the confound of stimulation intensity and significantly reduces visual effects (Amaya et al., 2023a). This allows us to test for the rhythmicity-dependent effects of FLS that drive differences in the subjective experience.

In this study, we implemented rhythmic 10 Hz FLS and frequency-matched arrhythmic FLS to test which thalamic nuclei display changes in functional connectivity during FLS-induced visual hallucinations. Additionally, we used a block design with rhythmic and arrhythmic flicker sequences to assess differential changes in whole-brain cortical activation, where we expected that rhythmic FLS would elicit stronger activation clusters in higher-order visual cortices compared to arrhythmic stimulation. To assess the resting-state functional connectivity, we utilised a fine anatomic parcellation of the thalamus (Rolls et al., 2020) and visual functional regions (Wang et al., 2015). We hypothesised there to be a rhythmicity-dependent effect of 10 Hz FLS on connectivity changes between higher-order visual cortices and thalamic nuclei, primarily anterior, ventral and mediodorsal nuclei, as found previously (Amaya et al., 2023b). The stronger activation and connectivity patterns during rhythmic 10 Hz FLS compared to arrhythmic would therefore be associated with differences in visual hallucination intensity.

## Methods

### Participants

Twenty participants took part in the experiment (fifteen male, five female; mean age ± standard deviation = 29 ± 7.9 years). Participants were recruited through word-of-mouth and student mailing lists within the Freie Universität Berlin. Eighteen participants were right-handed and two were left-handed according to the Edinburgh Handedness Inventory (Oldfield, 1971) (mean laterality quotient for right-handers (M ± SD) = 64.7 ± 18.7; laterality quotient for left-handers = -60; -20). Inclusion criteria involved no acute mental disorders, no current consumption of psychotropic medication and previous experience with FLS to minimise risk of aversive effects. Participants gave their written consent before proceeding with the experiment. The experimental design and materials were approved by the ethics committee at Freie Universität Berlin (application reference: 085/2021).

### Experimental materials and set-up

A stroboscope lamp with twelve 4500k J2 6V LEDs was used to generate the light stimulation, capable of delivering a maximum of 10,360 Lumens (Lumenate Growth Ltd., Bristol, United Kingdom). For study implementation, the lamp was programmed to deliver 8,125 Lumens, which is approximately 80% of its total capacity. The lamp was placed outside the scanner, perpendicular to the direction of the scanner bore (in place of the usual projector) and reflected by the MRI mirror system into the scanner bore. Convex lenses were placed above participants’ eyes to refocus the light and increase comparability of the set-up to previous studies (Amaya et al., 2023a; Montgomery et al., 2023). The lamp was interfaced with an Arduino (v1.8.16) to deliver FLS at two rhythmicity levels, namely: Rhythmic 10 Hz, which consisted of periodic light stimulation following a 0.3 duty cycle (30ms on, 70ms off), and Arrhythmic flicker [Figure 1A]. The arrhythmic flicker sequence used pairs of high frequency flashes embedded within longer intervals sampled from an exponential probability distribution (mean = 70 ms; see Arrhythmic_pairs_ condition from (Amaya et al., 2023a) for more details on arrhythmic sequence). Rhythmic 10 Hz FLS was applied to induce the highest intensity of visual hallucinatory phenomena, while arrhythmic stimulation elicits significantly fewer hallucinatory effects (Amaya et al., 2023a).

**Figure 1.**
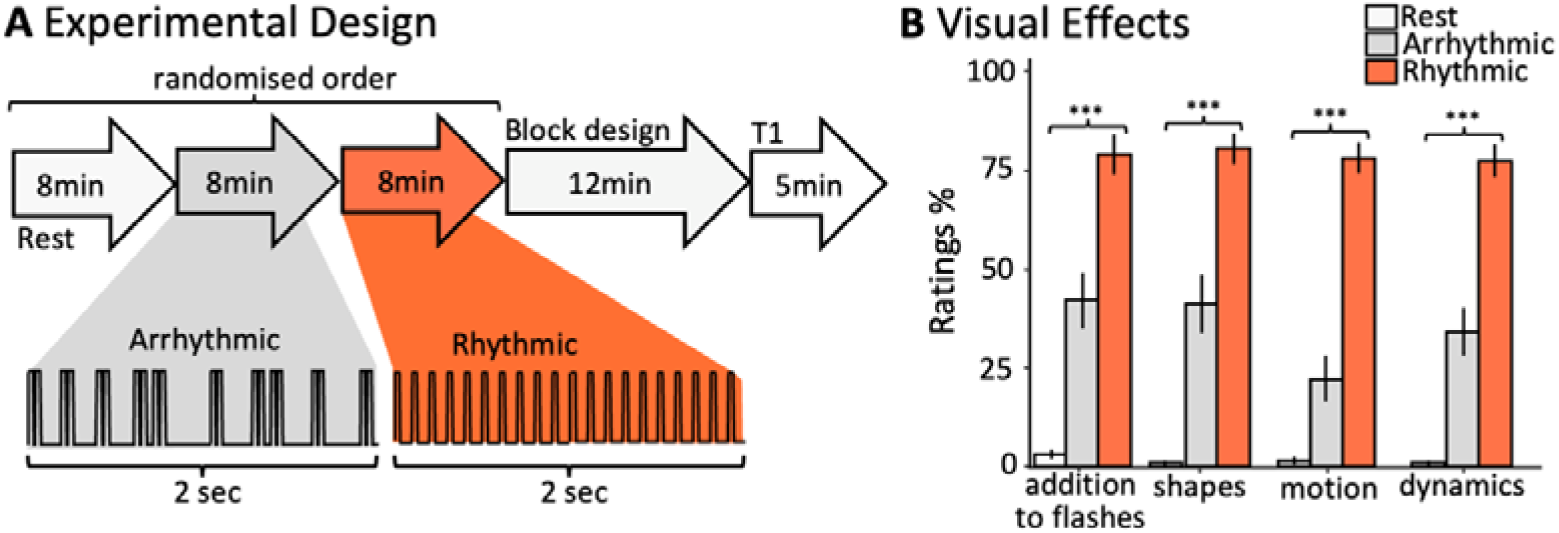
(A) The scanning session comprised of three eight-minute resting-state scans: (1) closed-eye rest in darkness (2) closed-eye rest with Arrhythmic FLS and (3) closed-eye rest with 10 Hz Rhythmic FLS, presented in a fully randomised order. After each resting-state scan, participants responded to four questions about their subjective experiences via the scanner intercom system. Thereafter, participants underwent a 12-minute run with eyes closed where rhythmic and arrhythmic FLS sequences were presented in a block design (see Methods for details). Each block lasted 20 seconds followed by a 20 second baseline interval. An anatomical scan was performed before participants were released from the scanner. (B) Mean scores of questionnaire items for each experimental condition, indicating that flicker rhythmicity significantly affects the reported intensity of phenomena additional to flashes, perceived shapes, motion, and experience dynamics. Stars represent the significance of a one-way repeated-measures ANOVA to test the effect of experimental condition on subjective ratings (*p* < 0.001 marked with ***). Error bars represent the standard error of means.

### Study design and procedure

MRI scanning was conducted at the Center for Cognitive Neuroscience Berlin (CCNB), Freie Universität Berlin. The scanning session comprised of three eight-minute resting-state scans: (1) closed-eye rest in darkness (2) closed-eye rest with 10 Hz Rhythmic FLS and (3) closed-eye rest with 10 Hz Arrhythmic FLS, presented in a fully randomised order [Figure 1C]. After each resting-state scan, the participants rated four questions (see below) about their subjective experiences via the scanner intercom system. Thereafter, participants underwent a 12-minute functional scan with eyes closed during which a block design with six conditions was applied: (1) darkness, (2) constant light, (3) 3 Hz Rhythmic FLS, (4) 3 Hz Arrhythmic FLS, (5) 10 Hz Rhythmic FLS, and (6) 10 Hz Arrhythmic FLS. Each block lasted 20 seconds followed by a 20 second baseline interval. Each condition was presented three times within three blocks of all six conditions, where within each block the order of conditions was randomized. Subsequently, a 5-minute anatomical scan was performed before participants were released from the scanner and experiment.

### FLS-induced phenomenology

Phenomenological aspects of FLS-induced effects were retrospectively assessed using four questionnaire items from an adapted version of the Stroboscopic Visual Experience Survey (SVES), which were previously identified as most characteristic of the subjective experience (Amaya et al., 2023a). The items were as follows: “Did your visual experience consist of anything in addition to blackness and/or flashes?”, “Did your visual experience consist of geometric formations including points, lines, shapes and/or patterns?”, “Did your visual experience include motion? Such as patterns or shapes that moved across, around or within your visual field?” and “Did your visual experience continuously change or evolve over time?”. Participants were asked to give an integer rating between 0 (no, not at all) and 100 (yes, very much).

### fMRI scanning

Participants were scanned using a 3T MAGNETOM Prisma Fit MRI scanner equipped with a 64-channel head coil (Siemens Healthineers, Erlangen, Germany). Resting-state fMRI images were acquired using a T2*-weighted echo planar imaging (EPI) sequence (48 axial slices acquired interleaved with multi-slice factor 3, in-plane resolution is 2.5 × 2.5 mm, slice thickness = 2.5 mm, flip angle = 70°, TR = 1500 ms, TE = 33 ms). To acquire structural images, a T1-weighted MPRAGE sequence was used (TR = 1930 ms, TE = 3.52 ms, flip angle = 8°, voxel size 0.8 × 0.8 × 0.8 mm). Cushioned head supports were fitted to minimise head motion.

### MRI data pre-processing

Data were pre-processed using SPM12 (www.fil.ion.ucl.ac.uk/spm/). Slice time correction and realignment was applied to the functional data before spatial normalisation to MNI152 space using unified segmentation in SPM12, which included reslicing to an isometric 2 mm voxel size (Ashburner and Friston, 2005).

### Cortical activation analysis

The realigned and normalised data was smoothed with an 8 mm FWHM Gaussian kernel. Statistical analysis was performed according to a standard general linear model (GLM) approach. The first-level design was specified to model the six task-conditions (Rhythmic 10 Hz, Arrhythmic 10 Hz, Rhythmic 3 Hz, Arrhythmic 3 Hz, Constant light, and No light) as independent regressors and motion parameters were modelled in six regressors of no interest. On the first level, we computed contrasts against implicit baseline for Rhythmic 10 Hz, Arrhythmic 10 Hz, Rhythmic 3 Hz, Arrhythmic 3 Hz, Constant light and No light conditions. These contrasts were entered into a second-level flexible factorial design, as implemented in SPM12, which included modelling of the subject factor (Gläscher and Gitelman, 2008). The second-level contrasts ‘10 Hz Rhythmic’ > ‘Implicit Baseline’ and ‘10 Hz Rhythmic’ > ‘10 Hz Arrhythmic’ were computed. All activation differences are reported at *p* < 0.05, family wise error (FWE) corrected on the cluster level, unless stated differently. Thresholded statistical parametric maps were rendered on a standard 3D brain template using MRIcroGL v1.2.20220720b (https://www.nitrc.org/projects/mricrogl/). Unthresholded t-maps can be found at https://neurovault.org/collections/FEKKFQRX/. All coordinates are reported in the MNI space of SPM12.

### Resting-state fMRI analysis

A low-pass Butterworth filter of 0.15 Hz was applied to realignment parameters to filter out factitious motion induced by magnetic field perturbations during respiration (Fair et al., 2020; Power et al., 2019; Gratton et al., 2020), before calculating the frame-wise displacement (FD) using BRAMILA tools (Power et al., 2012). Volumes that exceeded a threshold of 0.4 mm were masked during following analysis steps (*“scrubbing”*). The mean FD across scans and participants was 0.08 ± 0.04 (M ± SD) and the mean percentage of volumes scrubbed for each scan was 0.44 ± 0.67 (M ± SD), with the maximum percentage being 3.13%. Principal component analysis (CompCor) was performed using the DPABI toolbox (toolbox for Data Processing & Analysis of Brain Imaging, http://rfmri.org/dpabi) within the CSF/white matter mask to estimate nuisance signals (Behzadi et al., 2007). Anatomical masks for CSF, white and grey matter were derived from tissue-probability maps provided in SPM12. Smoothing was performed with a 3 mm FWHM Gaussian kernel to retain high spatial specificity for small regions of interest (Mikl et al., 2008; Pajula and Tohka, 2014; Chen and Calhoun, 2018). The first five principal components of the CompCor analysis, six head motion parameters, linear and quadratic trends as well as the global signal were used as nuisance signals to regress out associated variance. All analyses were additionally run without global signal regression (GSR) and included in the Supplement. Finally, the toolbox REST (www.restfmri.net) was used for temporal band-pass filtering (0.01-0.08Hz).

Anatomical regions of interest (ROIs) were extracted using the automated anatomical labelling atlas (AAL3; (Rolls et al., 2020)) for the thalamus and a volume-based maximum probability map (MPM) of visual topography (Wang et al., 2015) for cortical areas. Using probability maps of V1 and V2, overlapping ROIs at the midline were resolved by assigning voxels to the region with highest probability. Of thalamic ROIs, the Reuniens nucleus is only 8mm^3^ and was excluded from our analyses. Additionally, when the MPM was resliced to 2 mm^3^, hMST and IPS5 were no longer present in one hemisphere. For consistency, these ROIs were removed bilaterally.

For each ROI, mean BOLD time courses were extracted and temporal ROI-to-ROI Pearson correlations were calculated and subsequently Fisher z-transformed. We took the mean of correlation coefficients for ipsilateral and contralateral connections to give one bilateral functional correlation coefficient for each pair of ROIs. For all ROI-to-ROI pairs, we took the group-level average of correlation coefficients for closed-eye rest, Rhythmic and Arrhythmic resting-state scans and then computed the differences via matrix subtraction. A mask was applied so that only significant differences (paired t-test *p* < 0.05 FDR-corrected with the Benjamini-Hochberg method (Benjamini and Hochberg, 1995)) were shown. Baseline connectivity and unthresholded connectivity change matrices can be found in the Supplement (Figure S1-6).

### Linear mixed modelling

To test the relationship between subjective experience and connectivity changes, we ran linear mixed effect models with subjective ratings as a fixed effect and ROI-to-ROI functional connectivity as the dependent variable. Participants were included as random effects, and the models were fit using the restricted maximum likelihood method. Subjective ratings were taken as the average of the four questionnaire items for each participant for each condition. Akaike Information Criterion values were similar for models with random intercepts only and models with random intercepts and slopes, therefore models were run with random intercepts and slopes to account better for inter-subject variability. Additionally, we ran linear mixed models using subjective ratings as the fixed effect and the percent signal change of peak voxels as the dependent variable. Percent signal changes were extracted from peak voxels in the clusters identified by the 10 Hz Rhythmic > 10 Hz Arrhythmic contrast using the RFXplot toolbox implemented within SPM12 (Gläscher, 2009). To test the underlying assumptions of linear mixed modelling, Normal Q-Q plots were used to assess whether residuals of the linear mixed model fit followed a Gaussian distribution. Homoscedasticity was confirmed by plotting the residuals against predicted values and observing no pattern in the noise distribution. The assumption of independence was addressed by including participants as a random effect in the model.

## Results

### Subjective ratings

Repeated-measures ANOVAs identify a significant effect of condition on ratings of “Did your visual experience consist of anything in addition to blackness and/or flashes?” (F(2, 38) = 88.94, *p* < 0.001), “Did your visual experience consist of geometric formations including points, lines, shapes and/or patterns?” (F(1.5, 28.0) = 91.43, *p* < 0.001), “Did your visual experience include motion? Such as patterns or shapes that moved across, around or within your visual field?” (F(2, 38) = 122.93, *p* < 0.001) and “Did your visual experience continuously change or evolve over time?” (F(2, 38) = 85.11, *p* < 0.001), where all conditions produced ratings that were significantly different from each other for each item, with 10 Hz Rhythmic FLS generating the highest ratings followed by Arrhythmic and closed-eye rest [Figure 1B]. As ratings were not normally distributed (e.g., tendency for positive skew during closed-eye rest conditions), we additionally confirmed these results using repeated-measures Friedman non-parametric tests to test effects of experimental condition on perceived motion (X2(2) = 38, *p* < 0.001), dynamics (X2(2) = 34.2, *p* < 0.001), perceived patterns and shapes (X2(2) = 36.1, *p* < 0.001) and seeing something additional to flashes (X2(2) = 35.6, *p* < 0.001).

### Flicker-induced cortical activation

To reveal the cortical activation induced by FLS condition, we computed the second-level contrast 10 Hz Rhythmic > Implicit Baseline and 10 Hz Rhythmic > Arrhythmic FLS. The Rhythmic > Implicit Baseline cluster revealed increased activity in the entire occipital cortex, as shown in Figure 2A, with *p* < 0.05 FWE corrected on the cluster level. The activation cluster (K_E_=17,507) extends bilaterally across the occipital cortex (peak: x = 0, y = -82, z= 4, t-value = 11.87; subpeaks: x = 10, y = -76, z = 10, t-value = 11.05; x = 10, y = -88, z = 4, t-value = 10.46) with peaks in medial ventral occipital areas. Additionally, one cluster was revealed in the intraparietal area (peak: x = 24, y = -48, z = 46, t-value = 6.19, K_E_ = 387). On the peak level, there were small clusters in bilateral LGN regions of the AAL3 atlas (right peak: x = 26, y = -24, z = -2, t-value = 7.09; K_E_ = 45; left peak: x = 26, y = -24, z = -2, t-value = 7.77, K_E_ = 62). Meanwhile, the Rhythmic > Arrhythmic contrast unveiled a specific increase in activation of bilateral higher-order visual cortices (left cluster K_E_ = 691; right cluster K_E_ = 564), with *p* < 0.05 FWE corrected on the cluster level (left peak: x = -34, y = -80, z = 10, t-value = 4.21; left subpeaks: x = -32, y = -78, z = 4, t-value = 4.14; x = -28, y = -86, z = -10, t-value = 4.06; right peak: x = 32, y = -76, z = 12, t-value = 4.29; right subpeaks: x = 24, y = -70, z = 46, t-value = 3.84; x = 28, y = -72, z = 34, t-value = 3.78). The left peak is in LO2, and the right peak is not within a region of the MPM, however has a 12% probability of being part of LO2. One right subpeak (x = 24, y = -70, z = 46) is part of the IPS0 region of the MPM [Figure 2D]. Unthresholded t-maps, accessible via https://neurovault.org/collections/FEKKFQRX/, show relatively symmetrical bilateral activation. Additionally, 10 Hz Arrhythmic > Implicit Baseline, 10 Hz Rhythmic > 3 Hz Rhythmic and 3 Hz Rhythmic > 3 Hz Arrhythmic contrasts were computed and included in the Supplement (Figure S7).

**Figure 2.**
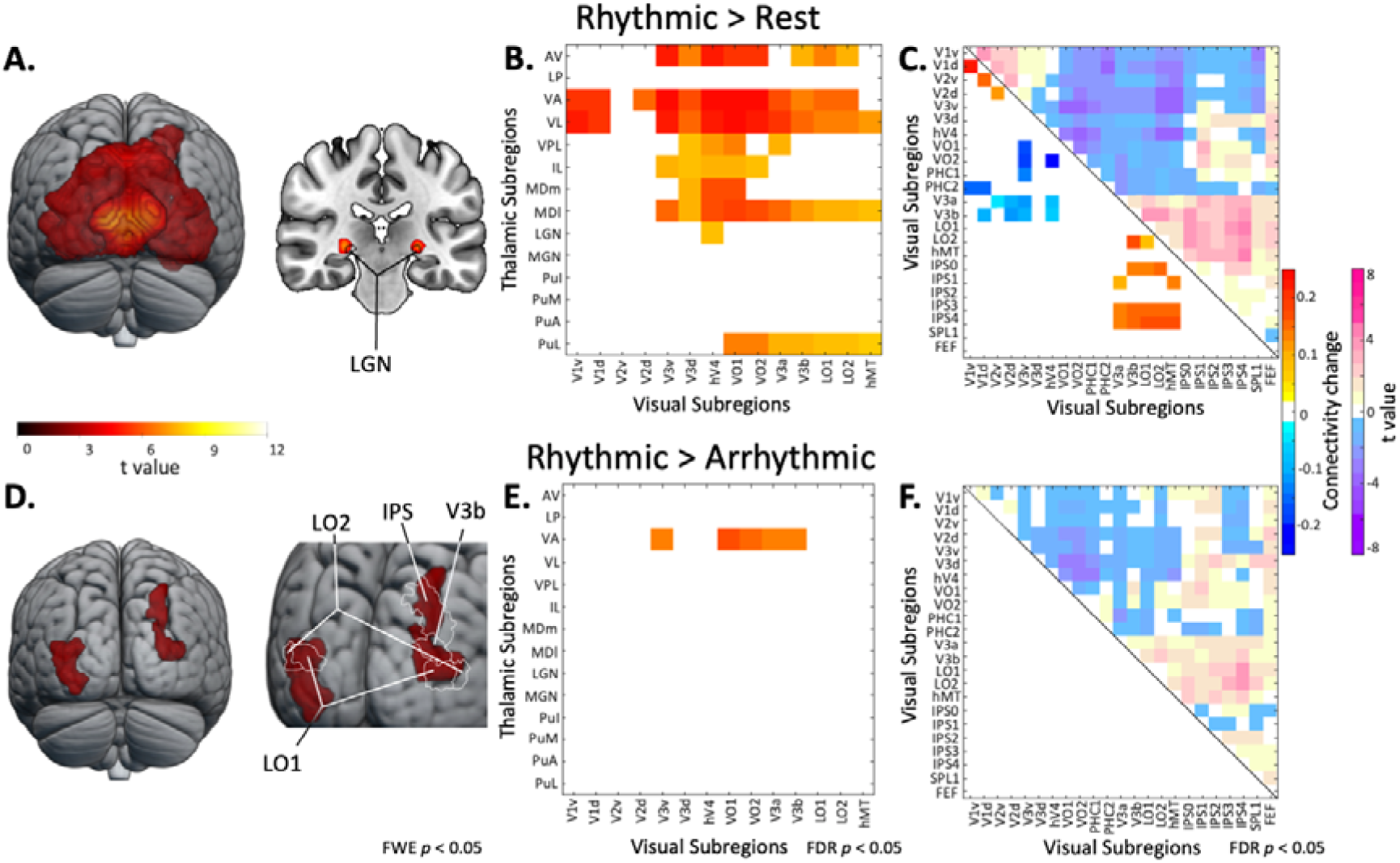
(A) Cortical activation clusters for Rhythmic 10 Hz against implicit baseline contrast (FWE *p* < 0.05 on the cluster level) reveals increased activation across the occipital cortex. When FWE *p* < 0.05 corrected on the peak level, the Rhythmic 10 Hz against implicit baseline contrast additionally reveals bilateral LGN activation. (B) Thalamocortical functional connectivity matrix showing the contrast of Rhythmic 10 Hz FLS against closed-eye rest reveals an increase in connectivity of anterior, ventral, mediodorsal and lateral pulvinar nuclei with visual cortices, which replicates previous findings (Amaya et al., 2023b). Unthresholded connectivity matrices are presented in the Supplement. (C) Corticocortical functional connectivity matrix showing the contrast of Rhythmic 10 Hz FLS against closed-eye rest reveals two clusters of hyperconnectivity within early visual cortices as well as between IPS and higher-order visual areas (e.g., LO1, V3a). There is an additional cluster of hypoconnectivity between early visual cortices and higher-order visual areas. (D) Rhythmic 10 Hz against Arrhythmic contrast reveals that the activation dependent on flicker rhythmicity is specific to higher-order visual cortices. (E) Thalamocortical functional connectivity matrix showing the contrast of Rhythmic 10 Hz FLS against Arrhythmic FLS. Connectivity between ventroanterior thalamic nuclei and higher-order visual areas (e.g., VO1) is significantly higher during Rhythmic FLS. (F) Corticocortical functional connectivity matrix of the contrast Rhythmic 10 Hz FLS against Arrhythmic FLS shows no significant differences in connectivity. All connectivity matrices are thresholded at *p* < 0.05 with FDR correction.

### Flicker-induced functional connectivity

We plotted connectivity matrices with 14 thalamic ROIs from the AAL3 atlas (Rolls et al., 2020) and 14 cortical ROIs from a maximum probability map of visual topography (Wang et al., 2015). The Rhythmic > Rest contrast shows an increase in connectivity between visual cortices and anterior, ventral and mediodorsal nuclei [Figure 2B], which replicates previous findings (Amaya et al., 2023b). Please note that connectivity between LGN and visual cortices was also increased during Rhythmic FLS, however this did not survive FDR correction due to existing baseline connectivity. Unthresholded connectivity matrices, which also include additional ROIs (e.g., intraparietal sulci), can be found in the Supplement (Figure S4-7). Figure 2C displays corticocortical functional connectivity matrices of Rhythmic FLS against closed-eye rest in darkness (*p* < 0.05 FDR-corrected) where there are two clusters of hyperconnectivity within early visual cortices and between IPS and higher-order visual areas (e.g., LO1, V3a) and an additional cluster of hypoconnectivity between early visual cortices and higher-order visual areas. This replicates findings from a previous dataset (Amaya et al., 2023b). When testing the Rhythmic against the Arrhythmic condition, we found a specific increase in connectivity (*p* < 0.05 FDR-corrected) between ventroanterior and higher-order visual cortices (e.g., VO1) during Rhythmic FLS [Figure 2E]. We found no significant differences in corticocortical connectivity changes between Rhythmic and Arrhythmic FLS [Figure 2F].

### Linear mixed modelling

We ran linear mixed models with subjective ratings of hallucination intensity as a fixed effect to determine if the reported intensity of visual effects (i.e., mean of four questionnaire items; see Methods) could predict changes in functional connectivity between thalamocortical ROI-to-ROI pairs. We applied a Bonferroni-corrected significance threshold (0.05/198). We found that subjective ratings significantly predicted connectivity between ventroanterior nuclei and VO1 (*p* = 0.0001, beta = 0.004; CI = [0.0017, 0.0053]; R^2^ adj = 0.51) and V3a (*p* = 0.0002, beta = 0.002; CI = [0.0011, 0.0036]; R^2^ adj = 0.28). Additionally, subjective ratings significantly predicted connectivity between ventrolateral nuclei and hV4 (*p* = 0.00025, beta = 0.003; CI = [0.0015, 0.0047; R^2^ adj = 0.44), VO1 (*p* = 3.9e-0.5, beta = 0.004; CI = [0.0021, 0.0055]; R^2^ adj = 0.68), V3a (*p* = 1.5e-0.5, beta = 0.003; CI = [0.0017, 0.0041]; R^2^ adj = 0.0.36), V3b (*p* = 0.0001, beta = 0.002; CI = [0.0012, 0.0033]; R^2^ adj = 0.39). We also tested the relationship between subjective ratings and the percent signal change in the peak voxels revealed by the 10 Hz Rhythmic > 10 Hz Arrhythmic activation contrast. Here, we only found a minor association between subjective ratings and activation differences in the peak voxel within LO2 of the left hemisphere (*p* = 0.027) and the right hemispheric peak voxel (*p* = 0.035), of which neither survive correction for multiple comparisons. Together, this shows that the subjective intensity of hallucinatory effects is more associated with ventral thalamic connectivity to higher-order visual cortices than the amount of cortical activation.

## Discussion

In this study, we tested for thalamocortical connectivity changes related to the subjective intensity of visual hallucinations induced by FLS, i.e. we tested for effects on rhythmicity, which influences the intensity of subjective experiences. First, we tested the effects of 10 Hz flicker rhythmicity on cortical activation, where we found a stronger activation of higher-order visual cortices during 10 Hz rhythmic FLS compared to arrhythmic control. Furthermore, we found a specific rhythmicity-dependent connectivity change between ventroanterior thalamic nuclei and higher-order visual cortices, where rhythmic FLS induced a stronger increase in functional connectivity. Consistently, rhythmic FLS also led to higher ratings of hallucination intensity across all measured domains, replicating findings from a previous study using the same questionnaire items (Amaya et al., 2023a). Linear mixed modelling revealed a positive relationship between thalamocortical connectivity and the reported intensity of hallucinatory effects, while the relationship between subjective ratings and cortical activation did not survive correction for multiple comparisons. In sum, we show that increased connectivity between ventroanterior thalamic nuclei and higher-order visual cortices relate to the increased intensity of visual hallucinatory effects during 10 Hz rhythmic flicker stimulation.

### Rhythmicity-specific activation in visual cortices

When assessing differences in cortical activation, we found that rhythmic 10 Hz FLS led to stronger activation specifically in higher-order areas of the visual cortex compared to arrhythmic FLS. This supports results from a previous fMRI study (Ffytche, 2008), which administered 8 Hz FLS alongside two control conditions: FLS with matched frequency but low brightness and FLS where the flashes were presented in pairs. The activation during 8 Hz FLS against low brightness control covered the occipital cortex similarly to our rhythmic 10 Hz FLS against baseline contrast. As the stimulation frequencies (i.e., 8 Hz, 10 Hz) lie within the alpha frequency range, which is known to elicit the most hallucinatory effects (Allefeld et al., 2011; Amaya et al., 2023a), it is expected that the activation results between studies are comparable. The activation clusters revealed in contrasts against both control conditions were in higher-order visual areas, such as the V5 complex, occipitotemporal cortices and intraparietal sulci (Ffytche, 2008). However, it must be noted that these findings were derived from a fixed effect model with six subjects. In our study, we refine these results by using a larger sample and a statistical model that accounts for the subject factor. Moreover, we developed and utilised an improved control condition consisting of arrhythmic flashes (Amaya et al., 2023a) accompanied by subjective ratings. Therefore, we provide more robust evidence for the increased activation of higher-order visual areas during hallucination-inducing FLS.

The visual areas that showed stronger activation during rhythmic 10 Hz FLS compared to arrhythmic FLS included LO1 and LO2, which are selective for orientation and shape perception, respectively (Malach et al., 1995; Grill-Spector et al., 1998; Silson et al., 2013); V3b, which is motion sensitive (Barton and Brewer, 2017); and the right intraparietal sulcus, which is part of the dorsal attention network (Vossel et al., 2014) but has also been found to respond selectively to simple geometric forms (Sereno and Maunsell, 1998). Together, the basic visual features that activates these cortical regions make up the key features of elementary visual hallucinations, namely coloured geometric patterns that move around the visual field. Indeed, 10 Hz rhythmic FLS was shown to increase ratings of perceived motion, shapes and patterns, found here and shown extensively in previous research (Smythies, 1959, 1960; Allefeld et al., 2011; Schwartzman et al., 2019; Bartossek et al., 2021; Amaya et al., 2023b, 2023a; Montgomery et al., 2023), suggesting that the activation of these visual cortical areas may contribute to the hallucinatory perception of elementary features.

Computational modelling has suggested that elementary visual hallucinations arise from perturbations of cortical excitation and inhibition in the primary visual cortex (Ermentrout and Cowan, 1979; Rule et al., 2011; Ermentrout and Billock, 2018), which can occur when rhythmic visual stimulation interacts with ongoing intrinsic oscillations. This was recently supported by an empirical study finding that standing waves are generated in the mouse visual cortex in response to rhythmic FLS (Gulbinaite et al., 2023). However, while this framework focuses on activity within primary visual cortices, most theories of conscious phenomenal experience require interactions with higher-order cortices, such as local recurrent interactions (Northoff and Lamme, 2020; Seth and Bayne, 2022), feedback interactions across the visual hierarchy (Corlett et al., 2019), or long-range interactions with supramodal regions (e.g. Global neural workspace theory (Dehaene and Changeux, 2011; Mashour et al., 2020)). In line with this, our data captures an increased BOLD signal in higher-order visual cortices during rhythmic FLS, which is in line with their functional contribution to colour, motion, and shape perception (as described above), and confirms previous reports (Ffytche, 2008). In this regard, while the interplay of cortical excitation and inhibition within primary visual cortices may initiate subsequent neural and phenomenal effects, our data emphasise the involvement of higher-order visual cortices as a relevant correlate of the subjective experience.

### Thalamocortical connectivity with ventroanterior thalamic nuclei

We found that rhythmic FLS led to selective increases in connectivity between ventroanterior nuclei and higher-order visual cortices, which was also associated with the intensity of reported visual effects. This presents a novel finding that refines previous work, which had found that 10 Hz FLS increased thalamocortical connectivity with anterior, ventral and mediodorsal nuclei compared to 3 Hz FLS (Amaya et al., 2023b), by removing confounds of stimulation intensity resulting from differences in flicker frequency. As ventroanterior nuclei are regarded as higher-order thalamic nuclei (Hwang et al., 2017), it suggests that thalamocortical hyperconnectivity within hallucinatory experiences relates to a higher-order function of the thalamus, such as in the coordination and integration of cortical activity. This aligns with a recent theory emphasising the role of thalamocortical interactions with higher-order nuclei in the formation of conscious experiences (Aru et al., 2019). Furthermore, ventral nuclei (including ventroanterior and ventrolateral divisions) are the only thalamic nuclei found to display above average connectivity to the right lateral visual network (Kumar et al., 2022), which is involved in shape and motion perception (Smith et al., 2009), demonstrating the feasibility that increased interactions between ventral thalamic nuclei and higher-order visual cortices are involved in FLS-induced visual effects.

Hallucinatory experiences can be elicited through various methods; however, the underlying neural state may be similar regardless of induction technique. For this reason, comparing findings from research using different hallucination induction methods can shed light on the shared neural effects and reinforce their likely relevance within visual hallucinatory experiences. For example, one study found that LSD induced hyperconnectivity between ventral and pulvinar thalamic nuclei and the sensorimotor network (Pizzi et al., 2023), while another found LSD induced hyperconnectivity between ventral thalamic nuclei and auditory-sensory cortical networks (Avram et al., 2022). While psychedelic experiences involve additional phenomenal and physiological effects, the shared connectivity pattern with FLS effects is consistently increased cortical connectivity with ventral thalamic nuclei. This further supports that connectivity with ventral, and more specifically ventroanterior, thalamic nuclei may be most relevant in the formation of visual hallucinatory experiences.

### Limitations and future directions

It is known that FLS can also induce other phenomenal effects, such as complex visual imagery (e.g., hallucinatory perception of realistic scenes; (Schwartzman et al., 2019; Bartossek et al., 2021)), enhancing the emotional response to music (Montgomery et al., 2023) and altering the sense of self (Schwartzman et al., 2019). The underlying mechanism of these additional phenomenal effects are pivotal to the potential clinical applications of FLS. For example, alterations in the sense of self may be linked to hypoconnectivity across the default mode network, as found with psychedelic interventions (Müller et al., 2018; Gattuso et al., 2022). However, these complex phenomena tend to emerge during FLS sessions that were designed to be maximally immersive, which is usually achieved through varying flicker frequencies and brightness as done in recreational settings. Future studies could utilize FLS sequences that are optimised to induce immersive experiences in order to assess the neural correlates of these additional phenomenal qualities. This would improve the comparability of FLS to other states, such as psychedelic experiences, while also lending to the clinical potential of FLS.

It is well established that FLS in the alpha frequency range elicits the most intense visual hallucinatory effects (Smythies, 1960; Allefeld et al., 2011; Schwartzman et al., 2019; Amaya et al., 2023a). Here, we administered 10 Hz FLS to be representative of the alpha frequency, however recent research found that using rhythmic flicker targeting the individual alpha frequency can lead to stronger effects of entrainment and changes in functional connectivity (Jaeger et al., 2023). Therefore, it is possible that accounting for the inter-individual variation in responsivity to alpha stimulation could unravel a stronger subject-specific relationship between cortical activation, connectivity changes and the reported visual effects.

In this study, we propose that the activation and connectivity differences induced by rhythmic and arrhythmic FLS are linked to differences in phenomenal experience. However, it remains possible that the neural effects are driven solely by flicker rhythmicity without being tied to phenomenal experiences. Still, as evidence indicates that FLS rhythmicity is critical for the hallucinatory experience (Ffytche, 2008; Amaya et al., 2023a), likely due to neural entrainment driving the effects (Notbohm 2016,Notbohm 2016,Schwartzman 2019,Jaeger 2023), there is reason to suggest that flicker rhythmicity and the visual effects are bound to each other. With this in consideration, while we do not present a causative link, our utilisation of a highly controlled experimental induction of visual hallucinations offers, to our knowledge, the most direct link between specific thalamocortical interactions and subjective experiences of visual hallucinations currently available in the literature. Importantly, this can be used to inform further clinical research into the mechanisms of hallucinatory symptomology.

## Conclusion

In sum, we show that higher-order visual cortices are specifically activated during rhythmic 10 Hz FLS compared to arrhythmic control. Furthermore, the rhythmicity of 10 Hz FLS selectively elicits an increase in functional connectivity between ventroanterior thalamic nuclei and higher-order visual cortices. These connectivity and activation changes reflect the difference in subjective ratings of perceived motion, dynamics, and shapes during FLS. Our results suggest that the role of thalamocortical hyperconnectivity during hallucinatory experiences is linked to a higher-order function of the thalamus, such as via changes in the coordination of cortical interactions.

## Acknowledgements

We thank Lumenate Growth Ltd (Bristol, United Kingdom) for generously providing experimental hardware free of charge.

## Notes

### Competing Interest Statement

The authors have declared no competing interest.

## References

Allefeld C, Pütz P, Kastner K, Wackermann J (2011) Flicker-light induced visual phenomena: Frequency dependence and specificity of whole percepts and percept features. Conscious Cogn 20:1344–1362.

Amaya IA, Behrens N, Schwartzman DJ, Hewitt T, Schmidt TT (2023a) Effect of frequency and rhythmicity on flicker light-induced hallucinatory phenomena. Plos One 18:e0284271.

Amaya IA, Schmidt ME, Bartossek MT, Kemmerer J, Kirilina E, Nierhaus T, Schmidt TT (2023b) Flicker light stimulation induces thalamocortical hyperconnectivity with LGN and higher-order thalamic nuclei. Imaging Neurosci 1:1–20.

Anticevic A, Cole MW, Repovs G, Murray JD, Brumbaugh MS, Winkler AM, Savic A, Krystal JH, Pearlson GD, Glahn DC (2014a) Characterizing Thalamo-Cortical Disturbances in Schizophrenia and Bipolar Illness. Cereb Cortex 24:3116–3130.

Anticevic A, Yang G, Savic A, Murray JD, Cole MW, Repovs G, Pearlson GD, Glahn DC (2014b) Mediodorsal and Visual Thalamic Connectivity Differ in Schizophrenia and Bipolar Disorder With and Without Psychosis History. Schizophrenia Bull 40:1227–1243.

Aru J, Suzuki M, Rutiku R, Larkum ME, Bachmann T (2019) Coupling the State and Contents of Consciousness. Front Syst Neurosci 13:43.

Ashburner J, Friston KJ (2005) Unified segmentation. NeuroImage 26:839–851.

Avram M, Brandl F, Knolle F, Cabello J, Leucht C, Scherr M, Mustafa M, Koutsouleris N, Leucht S, Ziegler S, Sorg C (2020) Aberrant striatal dopamine links topographically with cortico-thalamic dysconnectivity in schizophrenia. Brain 143:3495–3505.

Avram M, Müller F, Rogg H, Korda A, Andreou C, Holze F, Vizeli P, Ley L, Liechti ME, Borgwardt S (2022) Characterizing Thalamocortical (Dys)connectivity Following D-Amphetamine, LSD, and MDMA Administration. Biol Psychiatry: Cogn Neurosci Neuroimaging 7:885–894.

Barton B, Brewer AA (2017) Visual Field Map Clusters in High-Order Visual Processing: Organization of V3A/V3B and a New Cloverleaf Cluster in the Posterior Superior Temporal Sulcus. Front Integr Neurosci 11:4.

Bartossek MT, Kemmerer J, Schmidt TT (2021) Altered states phenomena induced by visual flicker light stimulation. Plos One 16:e0253779.

Bedford P, Hauke DJ, Wang Z, Roth V, Nagy-Huber M, Holze F, Ley L, Vizeli P, Liechti ME, Borgwardt S, Müller F, Diaconescu AO (2023) The effect of lysergic acid diethylamide (LSD) on whole-brain functional and effective connectivity. Neuropsychopharmacology 48:1175–1183.

Behzadi Y, Restom K, Liau J, Liu TT (2007) A component based noise correction method (CompCor) for BOLD and perfusion based fMRI. NeuroImage 37:90–101.

Benjamini Y, Hochberg Y (1995) Controlling the False Discovery Rate: A Practical and Powerful Approach to Multiple Testing. J R Stat Soc: Ser B (Methodol) 57:289–300.

Chen Z, Calhoun V (2018) Effect of Spatial Smoothing on Task fMRI ICA and Functional Connectivity. Front Neurosci 12:15.

Corlett PR, Horga G, Fletcher PC, Alderson-Day B, Schmack K, Powers AR (2019) Hallucinations and Strong Priors. Trends Cogn Sci 23:114–127.

Dehaene S, Changeux J-P (2011) Experimental and Theoretical Approaches to Conscious Processing. Neuron 70:200–227.

Ermentrout BG, Billock VA (2018) Flicker-Induced Phosphenes. Encyclopedia of Computational Neuroscience.

Ermentrout GB, Cowan JD (1979) A mathematical theory of visual hallucination patterns. Biol Cybern 34:137–150.

Ferri J et al. (2018) Resting-state thalamic dysconnectivity in schizophrenia and relationships with symptoms. Psychol Med 48:2492–2499.

Ffytche DH (2008) The hodology of hallucinations. Cortex 44:1067–1083.

Gattuso JJ, Perkins D, Ruffell S, Lawrence AJ, Hoyer D, Jacobson LH, Timmermann C, Castle D, Rossell SL, Downey LA, Pagni BA, Galvão-Coelho NL, Nutt D, Sarris J (2022) Default Mode Network Modulation by Psychedelics: A Systematic Review. Int J Neuropsychopharmacol 26:155–188.

Gläscher J (2009) Visualization of Group Inference Data in Functional Neuroimaging. Neuroinformatics 7:73–82.

Gläscher J, Gitelman D (2008) Contrast weights in flexible factorial design with multiple groups of subjects.

Grill-Spector K, Kushnir T, Hendler T, Edelman S, Itzchak Y, Malach R (1998) A sequence of object-processing stages revealed by fMRI in the human occipital lobe. Hum Brain Mapp 6:316–328.

Gulbinaite R, Nazari M, Rule ME, Bermudez-Contreras EJ, Cohen MX, Heimel JA, Mohajerani MH (2023) Spatiotemporal resonance in mouse primary visual cortex. bioRxiv:2023.07.31.551212.

Halassa MM, Sherman SM (2019) Thalamocortical Circuit Motifs: A General Framework. Neuron 103:762–770.

Hirschfeld T, Prugger J, Majić T, Schmidt TT (2023) Dose-response relationships of LSD-induced subjective experiences in humans. Neuropsychopharmacology:1–10.

Hwang K, Bertolero MA, Liu WB, D’Esposito M (2017) The Human Thalamus Is an Integrative Hub for Functional Brain Networks. J Neurosci 37:5594–5607.

Jaeger C, Nuttall R, Zimmermann J, Dowsett J, Preibisch C, Sorg C, Wohlschlaeger A (2023) Targeted rhythmic visual stimulation at individual participants’ intrinsic alpha frequency causes selective increase of occipitoparietal BOLD-fMRI and EEG functional connectivity. NeuroImage 270:119981.

Jan T, Castillo J del (2012) Visual Hallucinations: Charles Bonnet Syndrome. West J Emerg Medicine Integrating Emerg Care Popul Heal 13:544–547.

Kumar VJ, Beckmann CF, Scheffler K, Grodd W (2022) Relay and higher-order thalamic nuclei show an intertwined functional association with cortical-networks. Commun Biology 5:1187.

Lieslehto J, Jääskeläinen E, Kiviniemi V, Haapea M, Jones PB, Murray GK, Veijola J, Dannlowski U, Grotegerd D, Meinert S, Hahn T, Ruef A, Isohanni M, Falkai P, Miettunen J, Dwyer DB, Koutsouleris N (2021) The progression of disorder-specific brain pattern expression in schizophrenia over 9 years. Npj Schizophrenia 7:32.

Luppi AI, Carhart-Harris RL, Roseman L, Pappas I, Menon DK, Stamatakis EA (2021) LSD alters dynamic integration and segregation in the human brain. Neuroimage 227:117653.

Malach R, Reppas JB, Benson RR, Kwong KK, Jiang H, Kennedy WA, Ledden PJ, Brady TJ, Rosen BR, Tootell RB (1995) Object-related activity revealed by functional magnetic resonance imaging in human occipital cortex. Proc National Acad Sci 92:8135–8139.

Mashour GA, Roelfsema P, Changeux J-P, Dehaene S (2020) Conscious Processing and the Global Neuronal Workspace Hypothesis. Neuron 105:776–798.

Mikl M, Mareček R, Hluštík P, Pavlicová M, Drastich A, Chlebus P, Brázdil M, Krupa P (2008) Effects of spatial smoothing on fMRI group inferences. Magn Reson Imaging 26:490–503.

Montgomery C, Amaya IA, Schmidt TT (2023) Flicker light stimulation enhances the emotional response to music: A comparison study to the effects of psychedelics. PsyArXiv.

Müller F, Dolder PC, Schmidt A, Liechti ME, Borgwardt S (2018) Altered network hub connectivity after acute LSD administration. Neuroimage Clin 18:694–701.

Müller F, Lenz C, Dolder P, Lang U, Schmidt A, Liechti M, Borgwardt S (2017) Increased thalamic resting-state connectivity as a core driver of LSD-induced hallucinations. Acta Psychiat Scand 136:648–657.

Northoff G, Lamme V (2020) Neural signs and mechanisms of consciousness: Is there a potential convergence of theories of consciousness in sight? Neurosci Biobehav Rev 118:568–587.

Notbohm A, Herrmann CS (2016) Flicker Regularity Is Crucial for Entrainment of Alpha Oscillations. Front Hum Neurosci 10:503.

Notbohm A, Kurths J, Herrmann CS (2016) Modification of Brain Oscillations via Rhythmic Light Stimulation Provides Evidence for Entrainment but Not for Superposition of Event-Related Responses. Front Hum Neurosci 10:10.

Ommen MM van, Laar T van, Cornelissen FW, Bruggeman R (2019) Visual hallucinations in psychosis. Psychiatry Res 280:112517.

Pajula J, Tohka J (2014) Effects of spatial smoothing on inter-subject correlation based analysis of FMRI. Magn Reson Imaging 32:1114–1124.

Pizzi SD, Chiacchiaretta P, Sestieri C, Ferretti A, Tullo MG, Penna SD, Martinotti G, Onofrj M, Roseman L, Timmermann C, Nutt DJ, Carhart-Harris RL, Sensi SL (2023) LSD-induced changes in the functional connectivity of distinct thalamic nuclei. NeuroImage 283:120414.

Power JD, Barnes KA, Snyder AZ, Schlaggar BL, Petersen SE (2012) Spurious but systematic correlations in functional connectivity MRI networks arise from subject motion. NeuroImage 59:2142–2154.

Preller KH, Razi A, Zeidman P, Stämpfli P, Friston KJ, Vollenweider FX (2019) Effective connectivity changes in LSD-induced altered states of consciousness in humans. Proc National Acad Sci 116:2743–2748.

Prugger J, Derdiyok E, Dinkelacker J, Costines C, Schmidt TT (2022) The Altered States Database: Psychometric data from a systematic literature review. Sci Data 9:720.

Rolls ET, Huang C-C, Lin C-P, Feng J, Joliot M (2020) Automated anatomical labelling atlas 3. Neuroimage 206:116189.

Rule M, Stoffregen M, Ermentrout B (2011) A Model for the Origin and Properties of Flicker-Induced Geometric Phosphenes. Plos Comput Biol 7:e1002158.

Schmid Y, Enzler F, Gasser P, Grouzmann E, Preller KH, Vollenweider FX, Brenneisen R, Müller F, Borgwardt S, Liechti ME (2015) Acute Effects of Lysergic Acid Diethylamide in Healthy Subjects. Biol Psychiat 78:544–553.

Schmidt TT, Berkemeyer H (2018) The Altered States Database: Psychometric Data of Altered States of Consciousness. Front Psychol 9:1028.

Schwartzman DJ, Schartner M, Ador BB, Simonelli F, Chang AY-C, Seth AK (2019) Increased spontaneous EEG signal diversity during stroboscopically-induced altered states of consciousness. Biorxiv:511766.

Sereno AB, Maunsell JHR (1998) Shape selectivity in primate lateral intraparietal cortex. Nature 395:500–503.

Seth AK, Bayne T (2022) Theories of consciousness. Nat Rev Neurosci 23:439–452.

Sherman SM, Guillery RW (2006) Exploring the thalamus and its role in cortical function, 2nd ed. MIT Press.

Shine JM, Lewis LD, Garrett DD, Hwang K (2023) The impact of the human thalamus on brain-wide information processing. Nat Rev Neurosci:1–15.

Silson EH, McKeefry DJ, Rodgers J, Gouws AD, Hymers M, Morland AB (2013) Specialized and independent processing of orientation and shape in visual field maps LO1 and LO2. Nat Neurosci 16:267–269.

Smith SM, Fox PT, Miller KL, Glahn DC, Fox PM, Mackay CE, Filippini N, Watkins KE, Toro R, Laird AR, Beckmann CF (2009) Correspondence of the brain’s functional architecture during activation and rest. Proc National Acad Sci 106:13040–13045.

Smythies JR (1959) The Stroboscopic Patterns. Br J Psychol 50:305–324.

Smythies JR (1960) The Stroboscopic Patterns. Brit J Psychol 51:247–255.

Vossel S, Geng JJ, Fink GR (2014) Dorsal and Ventral Attention Systems. Neurosci 20:150–159.

Wang L, Mruczek REB, Arcaro MJ, Kastner S (2015) Probabilistic Maps of Visual Topography in Human Cortex. Cereb Cortex 25:3911–3931.

