## Supplementary Information for "Thalamocortical interactions reflecting the intensity of flicker light-induced visual hallucinatory phenomena"

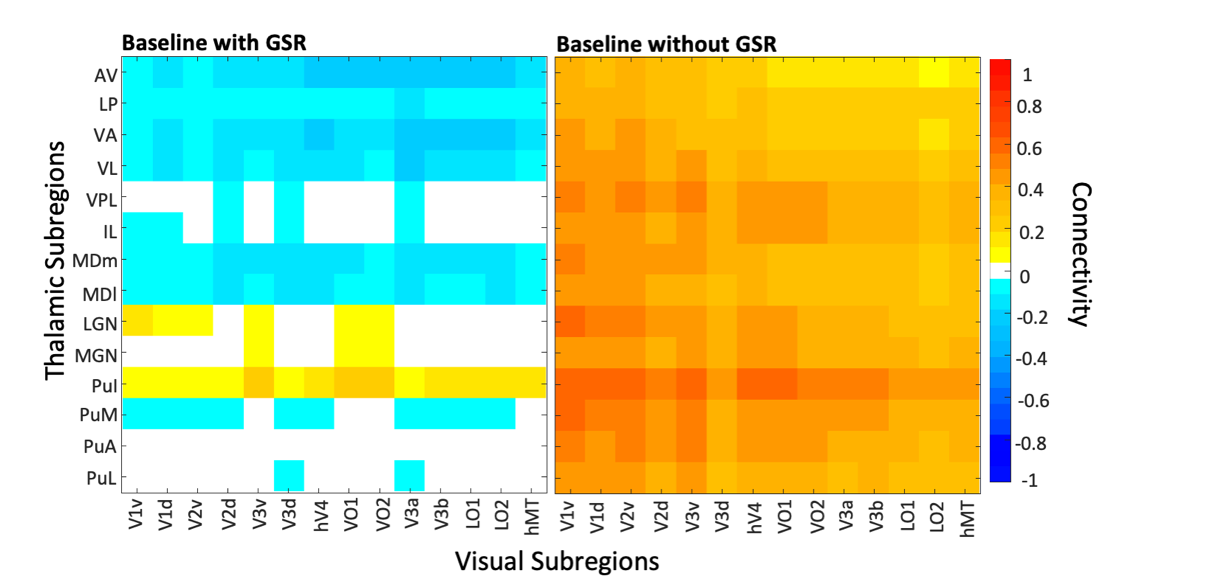

*Figure S1.* Connectivity matrices of thalamic and visual subregions during closed-eye rest in darkness. There is baseline connectivity between LGN and inferior pulvinar and visual cortices, which is expected due to their known involvement in visual processes. This likely also explains why FLS conditions were not shown to induce a significant (FDR-corrected *p* < 0.05) increase in LGN connectivity with visual cortices, as baseline correlations were already high. It is possible that some participants, although instructed to have eye closed, may have opened their eyes leading to higher correlations between LGN, pulvinar and visual cortices during baseline. Aside from this, the selective connectivity between LGN, inferior pulvinar and visual cortices during baseline measurements gives support to the specificity of the thalamic parcellation scheme. It is also evident that including global signal regression (GSR) during pre-processing steps introduces anti-correlations in some ROI-to-ROI pairs, which is a well-known artifact of GSR. However, reporting results using the contrast between conditions allows us to capture the relative change in connectivity, and therefore the introduction of anti-correlations does not bias the final reported results, as later demonstrated with Supplement Figure 3 and 4.

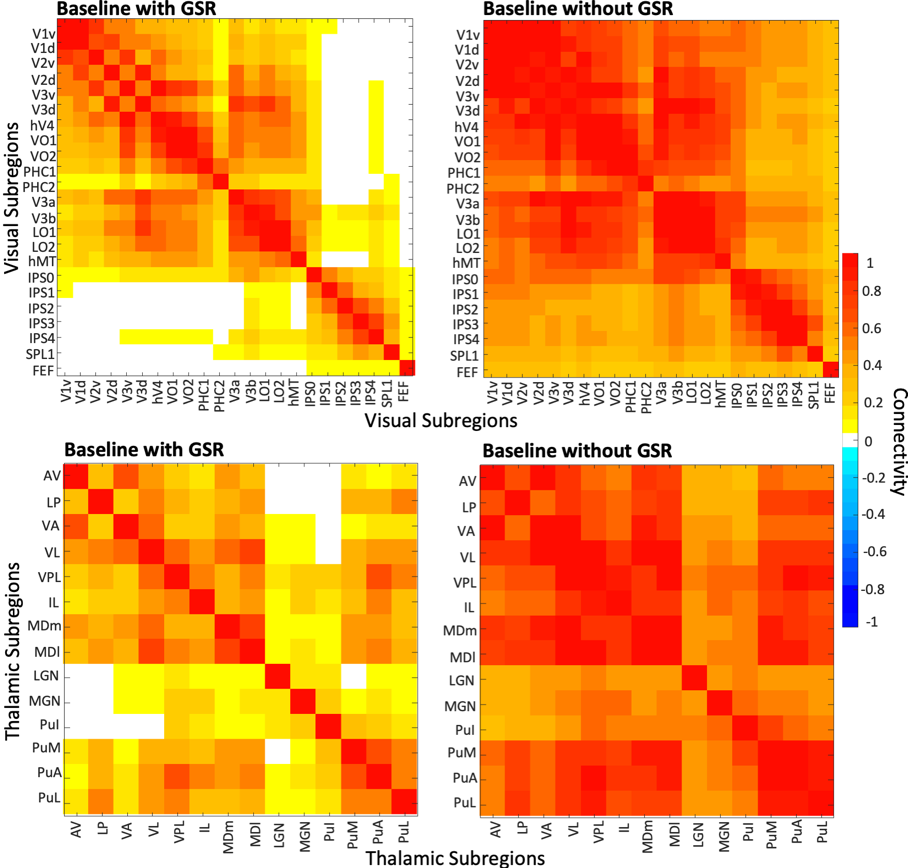

*Figure S2.* Connectivity matrices of interconnectivity between visual cortical regions (top) and thalamic subregions (bottom) during closed-eye rest in darkness. LGN has very weak baseline connectivity with ventral (i.e., ventroanterior and ventrolateral) thalamic nuclei, supporting that the results pinpointing strong connectivity changes for ventroanterior nuclei do not arise from a spatial smoothing effect from LGN signals. Correlations are lower when GSR is applied, however the overall interconnectivity patterns remain relatively consistent for both pre-processing lines.

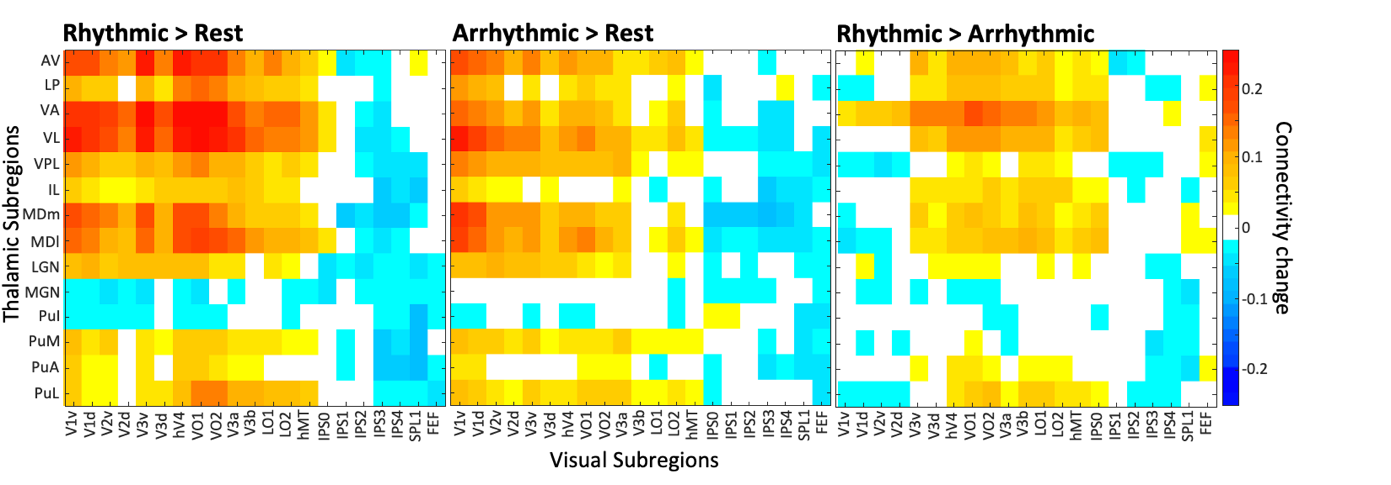

*Figure S3.* Unthresholded thalamocortical connectivity matrices for Rhythmic against Rest, Arrhythmic against Rest and Rhythmic against Arrhythmic contrasts with global signal regression applied. It is evident that there are increases in connectivity with visual cortices and LGN, anterior, ventral and mediodorsal nuclei. Due to baseline visual cortical connectivity with LGN, the increase induced by FLS does not survive conservation FDR correction for multiple comparisons. Additional connectivity effects seen for other nuclei, such as lateroposterior (LP) and ventroposterolateral (VPL) are likely due to effects of smoothing, as the nuclei have spatial proximity to ventral and mediodorsal nuclei that show stronger effects. There is also an increase in connectivity with the intraparietal sulcus (IPS0), which aligns with the activation cluster peak found in IPS0 in the Rhythmic > Arrhythmic activation contrast, however this would not survive correction for multiple comparisons.

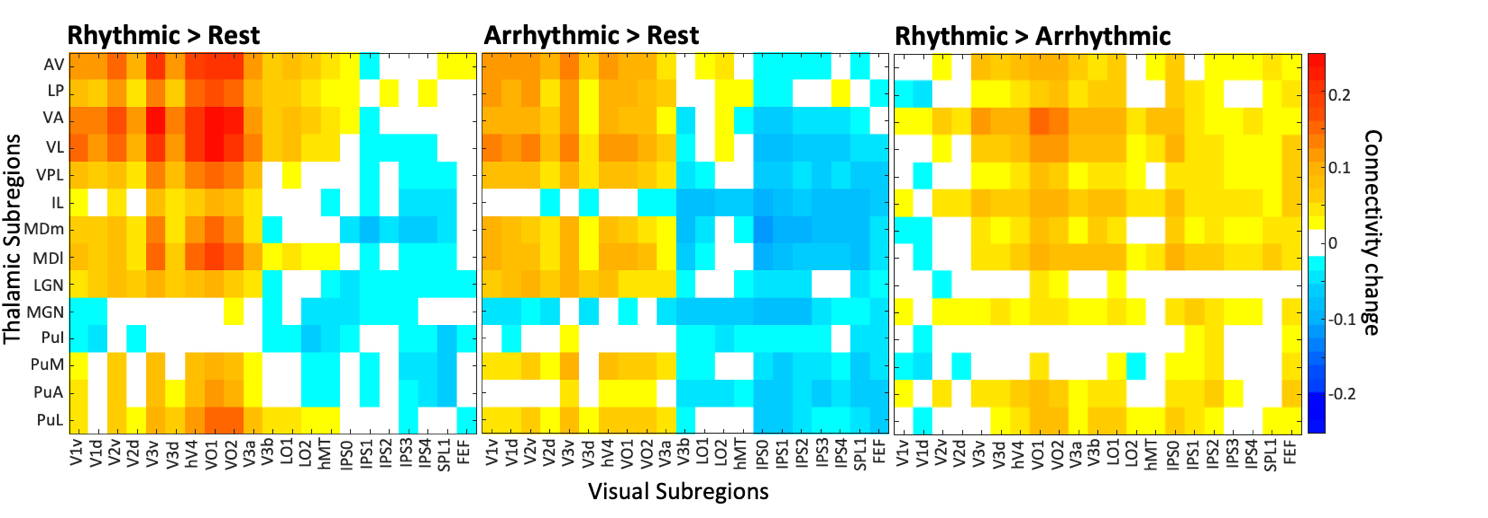

*Figure S4.* Unthresholded thalamocortical connectivity matrices for Rhythmic against Rest, Arrhythmic against Rest and Rhythmic against Arrhythmic contrasts when global signal regression (GSR) was not applied. Connectivity changes are highly similar as with GSR, indicating the stability and robustness of the main reported findings.

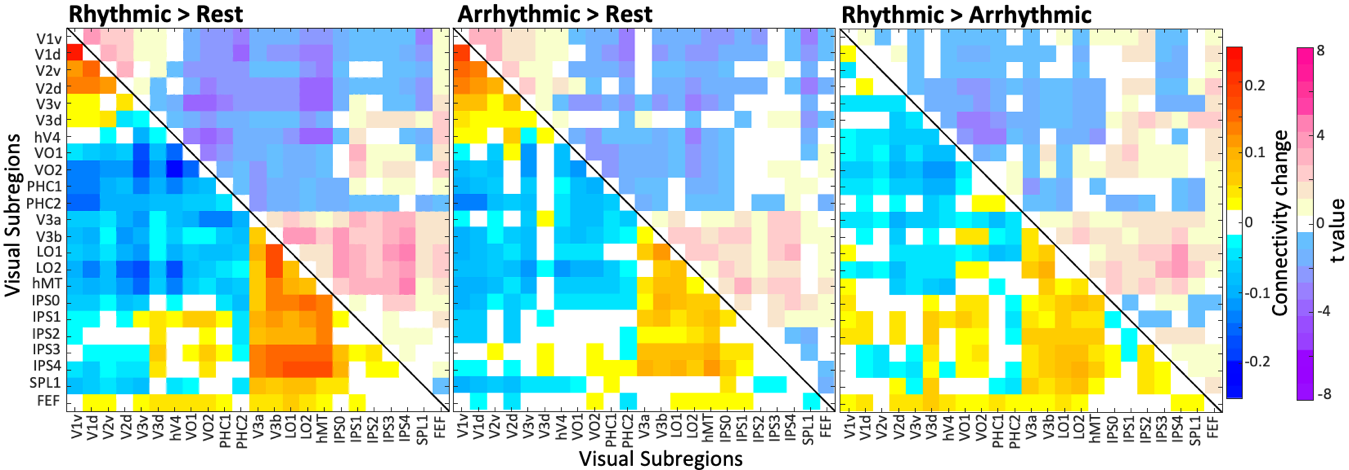

*Figure S5.* Unthresholded corticocortical connectivity matrices for Rhythmic against Rest, Arrhythmic against Rest and Rhythmic against Arrhythmic contrasts with global signal regression. The Rhythmic against Arrhythmic contrast shows slight changes in connectivity that are similar to the Rhythmic against Rest contrast, however the weaker effects mean that no connectivity changes survived FDR corrections.

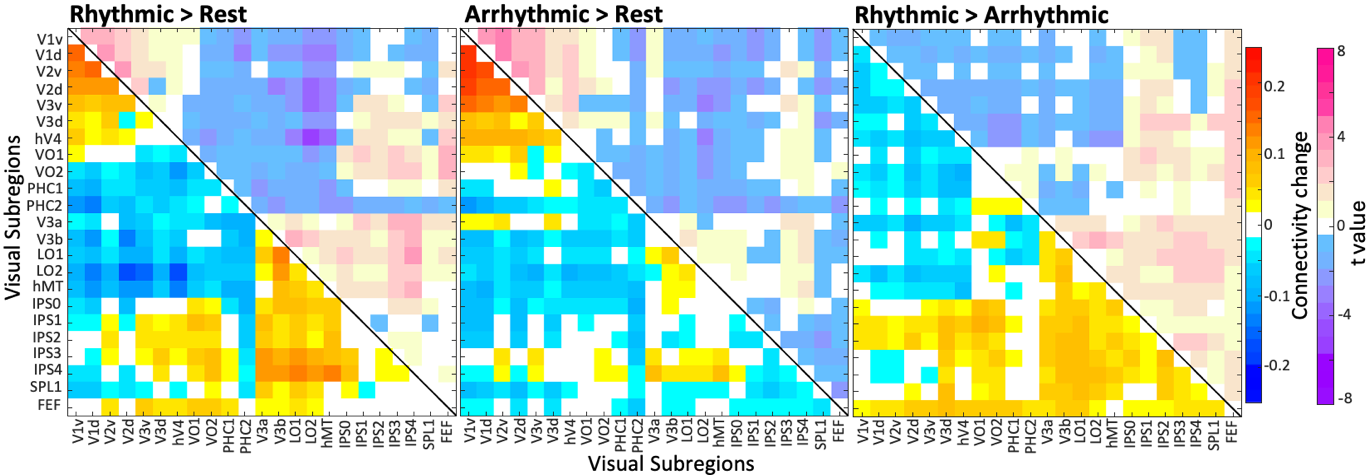

*Figure S6.* Unthresholded corticocortical connectivity matrices for Rhythmic against Rest contrast, Arrhythmic against Rest contrast and Rhythmic against Arrhythmic contrasts without global signal regression. The connectivity changes are highly similar as when GSR is applied, albeit slightly weaker. The similarity of the findings with and without GSR indicate that the findings are stable across variations in signal pre-processing.

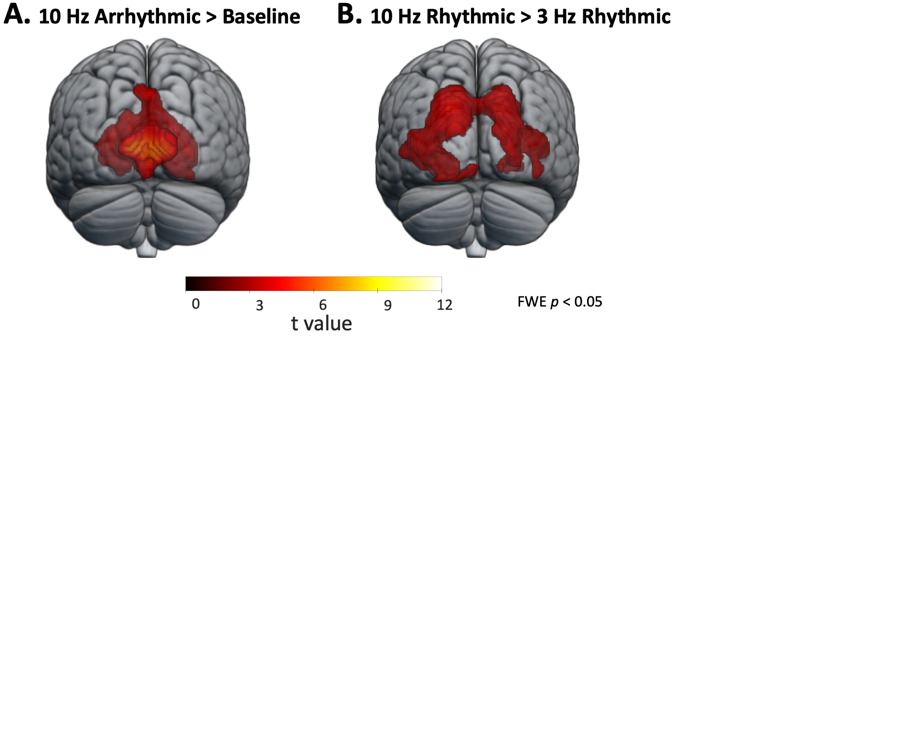

*Figure S7.* (A) Arrhythmic FLS > Implicit Baseline contrast (p < 0.05 FWE corrected on the cluster level) reveals increased activation of the ventral medial occipital areas akin to the early visual cortices. (B) 10 Hz Rhythmic > 3 Hz Rhythmic contrast (p < 0.05 FWE corrected on the cluster level) reveals large bilateral activation in higher-order visual cortices. While similar cortical areas are activated as in the 10 Hz Rhythmic > 10 Hz Arrhythmic contrast, the 10 Hz Rhythmic > 3 Hz Rhythmic contrast leads to a much larger activation cluster across various visual areas that likely arises from differences in stimulation intensity. However, as there are differences in visual hallucination intensity across flicker frequency, with 10 Hz induces more visual effects than 3 Hz, it supports that the activation of higher-order visual cortices likely corresponds to the subjective visual experience. The 3 Hz Rhythmic > 3 Hz Arrhythmic contrast reveals no significant activation clusters, which is also expected due to the frequency x rhythmicity interaction found previously (Amaya et al., 2023).
